# Exploring the diversity and spatial distribution of *Anopheles* mosquitoes and their associated *Plasmodium* species in Tanzania using the ANOSPP amplicon panel

**DOI:** 10.64898/2026.06.01.729474

**Authors:** Maneno E. Baravuga, Marilou M. Boddé, Alex Makunin, Praise Michael, Amos T. Mlalwe, Charles D. Mwalimu, Petra Korleviċ, Heather M. Ferguson, Henrik Salje, Mara K. N. Lawniczak, Nicodem J. Govella

## Abstract

Species-level identification of malaria vectors and parasites is essential for targeted vector control, surveillance, and clinical case management. However, routine surveillance often relies on morphology, species group-restricted PCR assays, and parasite-detection workflows optimised for *Plasmodium falciparum*, potentially masking wider vector and parasite diversity. To address this gap, we applied the previously published ANOSPP amplicon sequencing panel to *Anopheles* mosquitoes collected across 25 districts in mainland Tanzania between 2020 and 2023, using it to characterise vector and parasite diversity, evaluate routine species assignments, and examine how primary-vector occurrence related to national malaria endemicity patterns. Of 71,146 mosquitoes collected, 10,020 were morphologically identified as *Anopheles*. ANOSPP-based identification resolved 15 *Anopheles* taxa, including the first Tanzanian record of *Anopheles funestus*-like mosquitoes (n = 10) and the first resolution of *Anopheles longipalpis* to Type C in the country (n = 13). Using ANOSPP as the reference, morphology misclassified 6.2% of ANOSPP-analysed specimens, with *Anopheles rufipes* and *Anopheles maculipalpis* frequently misclassified, and *Anopheles pretoriensis* and *Anopheles marshallii* sensu lato completely missed. Five *Plasmodium* species were detected across 37 parasite-positive mosquitoes: *P. falciparum*, *P. vivax*, *P. ovale*, *P. malariae*, and *P. caprae*. *Plasmodium caprae* was detected only in *Anopheles arabiensis* (n = 6), representing the first record of this parasite in Tanzania and its first detection in *An. arabiensis* globally. Co-occurrence of *An. arabiensis*, *An. funestus* s.s., and *An. gambiae* s.s. was more common in higher malaria transmission strata, whereas occurrence of *An. arabiensis* alone predominated in very-low and low strata. These findings show that integrating ANOSPP into surveillance can substantially improve vector and parasite resolution, uncover overlooked diversity, and strengthen species-informed malaria surveillance.

**Author summary:** Effective malaria control requires accurate data on mosquito species distributions and the parasites they carry. However, in many settings surveillance still relies mainly on morphological identification or molecular tests targeting limited vector groups, while parasite detection often focuses on *Plasmodium falciparum*. This limits understanding of mosquito and parasite diversity, obscuring transmission dynamics. We used the previously published ANOSPP amplicon panel to analyse *Anopheles* mosquitoes across Tanzania. This revealed hidden biological complexity with direct implications for malaria transmission, resolving 15 mosquito and five *Plasmodium* species. Notably, for the first time in Tanzania, we detected the goat-associated parasite *Plasmodium caprae* in *Anopheles arabiensis* and identified *An. funestus*-like, whose vectorial capacity remains unclear, and *An. longipalpis* Type C, implicated in malaria transmission in dryland Kenya. Interestingly, co-occurrence of all three major Tanzanian vectors aligned with higher transmission settings, whereas areas where *An. arabiensis* was the sole vector were mainly in lower transmission settings. These findings show that species-resolved surveillance can reveal hidden biological complexity and provide a stronger basis for locally targeted malaria control.

## Introduction

Human malaria transmission in Tanzania involves a broader diversity of *Anopheles* vectors and *Plasmodium* parasites than routine entomological surveillance typically captures [1–3]. Although the primary vectors, *Anopheles gambiae* sensu stricto, *An. arabiensis*, and *An. funestus* sensu stricto, are well studied [1,4,5], other species have also been implicated in transmission. For example, *An. merus* has repeatedly been found infected with *Plasmodium* sporozoites in coastal areas, suggesting a focal role in malaria transmission [6–10], and several less-studied species, including *An. rivulorum, An. parensis, An. marshallii, An. leesoni, An. coustani, An. pharoensis,* and *An. squamosus*, have also been reported carrying *Plasmodium* sporozoites in the country [8,11–19]. However, these species have received little systematic follow-up and are rarely resolved to species level in routine entomological surveillance, where constraints in morphological expertise and PCR assays that target only selected species groups often lead them to be grouped as “other *Anopheles*” or reported only at complex or group level, obscuring their potential roles in transmission [20]. This is problematic because even closely related species can differ substantially in biting time, host preference, resting behaviour, ecological niche, and response to vector control interventions [21–30]. The parasite component of transmission is similarly incompletely captured. Recent surveys indicate that all major human *Plasmodium* species are present in Tanzania [2,31–34], yet their occurrence in vectors, and the mosquito species responsible for transmitting them, remain poorly characterized. As a result, the roles and spatial distributions of both *Anopheles* vectors and their associated *Plasmodium* species remain incompletely resolved.

Established vector surveillance systems in Tanzania are constrained by limited taxonomic resolution and uneven spatial coverage [35,36]. Approximately 50 *Anopheles* species have been reported in Tanzania, but most records rely solely on morphological keys, which have limited reliability for distinguishing sibling species without molecular validation [35,37,38]. Molecular diagnostics such as PCR assays have therefore been widely adopted and have improved identification, but they are designed for a limited set of species within the *An. gambiae* complex [39] and *An. funestus* group [40], leaving most other *Anopheles* unresolved at species level. These assays can also yield misleading results when initial morphological sorting is incorrect [20]. For example, standard *An. funestus* multiplex PCR can misidentify several non-*funestus* species as *An. leesoni* [41]. In addition, polymorphisms in primer-binding regions may cause non-amplification [11]. Consequently, both well-recognised primary vectors and routinely overlooked secondary taxa remain incompletely mapped and inconsistently monitored, creating persistent blind spots in national malaria surveillance [11,13,42]. These gaps have real epidemiological implications. In Madagascar, for example, *An. coustani* was long regarded as a minor vector but was later shown to contribute substantially to local malaria transmission [43–45]. This illustrates how neglecting secondary vectors can obscure transmission dynamics, misdirect control efforts, and perpetuate blind spots in elimination planning. Addressing this challenge requires surveillance approaches that move beyond morphology and single-locus PCR to achieve species-resolved detection across the full diversity of malaria vectors in the country.

Similar challenges persist in parasite surveillance. Although *P. falciparum* accounts for most severe malaria cases and malaria-related deaths, a fact that has understandably shaped surveillance priorities [46], this focus has also contributed to under-detection of other human and potentially zoonotic *Plasmodium* species. Within entomological programmes, parasite detection relies largely on circumsporozoite protein enzyme-linked immunosorbent assay (CSP-ELISA), the standard method for detecting sporozoites in mosquitoes [47–49]. Although CSP-ELISA can distinguish species when species-specific antibodies are used [48], many routine surveillance workflows, particularly in settings where *P. falciparum* predominates, are configured primarily for *P. falciparum* [1,46]. Expanding detection to additional species generally requires separate single-target assays, increasing workload, reagent use, and per-sample costs [47,50]. CSP-ELISA performance is also a concern: reports of false-positive results [51,52] and lower sensitivity relative to molecular assays [47,49,53] further indicate that sporozoite-rate estimates and vector incrimination may be unreliable without molecular confirmation. As a result, non-*falciparum* infections in vectors may remain undetected or incompletely characterized, obscuring epidemiologically relevant parasite diversity. This diversity matters because non-*falciparum* parasites differ substantially in their infection dynamics and public-health implications. *Plasmodium vivax* and *P. ovale* can relapse from dormant liver stages [54–56], and *P. malariae* can persist as chronic, often low-density blood-stage infections for years or even decades [34,57,58], while mixed-species infections may complicate diagnosis and treatment and have been associated with severe clinical outcomes [59,60]. Moreover, reports of zoonotic and sylvatic *Plasmodium* infections elsewhere show that animal-associated parasites can be missed by surveillance systems designed around a narrow set of human malaria species [61,62]. Without species-resolved detection, these infections remain poorly represented in surveillance data, limiting accurate burden estimation and weakening the correspondence between entomological indicators and the parasite species causing human disease.

Resolving these surveillance gaps at scale is increasingly important as countries such as Tanzania adopt subnational tailoring approaches to optimize malaria strategies and interventions according to local contexts [63,64]. Notably, the World Health Organisation (WHO) reference manual on subnational tailoring lists parasite species distribution and mosquito vector species among the priority indicators for subnational stratification [65]. In practice, however, operational stratification in Tanzania has so far been built primarily on infection-burden indicators such as parasite prevalence and case incidence [64], with species-specific vector and parasite data not yet routinely incorporated. This is problematic because malaria vector species differ in biting behaviour [66] and vectorial capacity [5,67] and respond differently to vector control interventions [29,30], while *Plasmodium* species differ in treatment requirements, relapse potential, persistence, and clinical implications [54,55,58–60]. For instance, previous work in Tanzania found that the spatial occurrence of *An. funestus* s.s. and *An. gambiae* s.s. was associated with areas of higher malaria transmission, despite their lower abundance relative to *An. arabiensis* [1]*. Anopheles funestus* s.s. has a higher vectorial capacity than *An. arabiensis* owing to its longer survival [67], strong human-feeding preference [24] and greater susceptibility to parasite infection [5]. This example shows that without species-resolved surveillance, it remains difficult to identify the biological drivers of local transmission and to determine how interventions can be optimized for maximum impact [28–30].

To address these challenges, this study employed the ANOSPP amplicon sequencing panel, a genomic tool capable of simultaneously identifying *Anopheles* species and *Plasmodium* species from the same mosquito specimen [68–70]. By using curated reference sequence databases, the panel provides species-level resolution across the genus *Anopheles* and discriminates multiple *Plasmodium* lineages with available reference sequences, while requiring only genus-level morphological identification and reducing reliance on detailed, error-prone morphological sorting. This study specifically (i) documents the diversity and geographical distribution of *Anopheles* mosquitoes and their associated *Plasmodium* species across Tanzania; (ii) compares ANOSPP-based identifications with conventional morphology to quantify misclassification and recovery of overlooked taxa; and (iii) assesses how the spatial composition of primary vectors relates to national malaria transmission strata as defined by the Tanzania National Malaria Control Program (NMCP) [64].

## Materials and methods

### Ethics statement

Ethical clearance for this study was obtained from the National Institute for Medical Research (NIMR/HQ/R.8a/Vol.IX/3390) and the Institutional Review Board of the Ifakara Health Institute (IHI-IRB No. 9-2020). Mosquito sampling was conducted in private households accessed with the permission of the household heads. Before participation, all volunteers received a detailed verbal explanation of the study objectives and procedures, followed by written informed consent. The consent form and accompanying information sheet emphasised confidentiality, voluntary participation, and the right to withdraw at any point without consequence, as well as potential risks and anticipated benefits. No protected or endangered species were sampled during the study. Permission to publish this study was granted by the National Institute for Medical Research, Tanzania (No. BD.242/437/01C/213).

### Research design

This study analysed entomological samples and data collected between December 2020 and December 2023 from 25 ecologically diverse districts across mainland Tanzania. As a large country with a significant malaria burden, and substantial ecological variability from coastal lowlands to the Great Rift Valley, Tanzania offers a representative landscape for investigating malaria vector-parasite dynamics. The selected districts are shown in Fig 1 and described in more detail in [36] and S1 Fig. These sites were part of the national malaria vector surveillance system coordinated by the National Malaria Control Programme’s Malaria Vector Surveillance (NMCP-MVES) [1], chosen to reflect a range of ecological zones, intervention strategies, malaria endemicity levels, and anthropological contexts. The study employed a rolling cross-sectional surveillance design [36]. Within each selected district, one of the three NMCP-designated sentinel villages was randomly chosen. In each village, three sub-villages were selected, and four households were enrolled per sub-village for entomological monitoring. Each sub-village was surveyed three times during the study period, with one night of mosquito sampling per visit. To enhance spatial coverage and minimize pseudo-replication, new households were selected for each round. Seasonal variation was addressed by ensuring that each village was sampled at least once during both wet and dry seasons [36].

**Fig 1.**
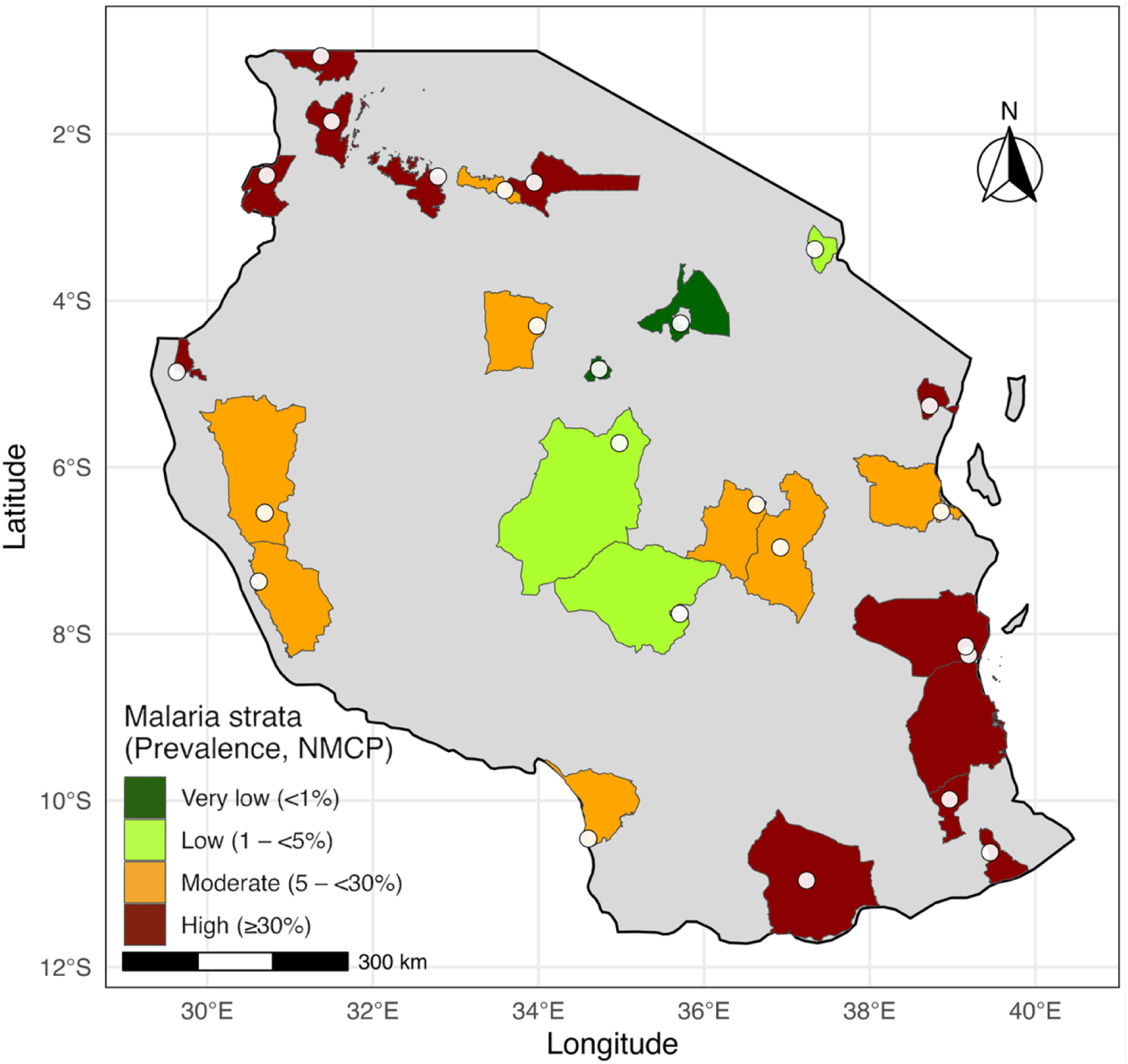
Study area. Map of 25 districts sampled across mainland Tanzania, coloured by National Malaria Control Programme transmission strata (Very low, Low, Moderate, High) [64]. White dots indicate sampling sites. District boundaries are based on the national administrative level-2 shapefile; however, the boundaries of some districts were enlarged for visual clarity.

### Mosquito collection

Mosquito collections were conducted using a combination of mosquito trapping methods, namely: (i) Mosquito Electrocuting Traps (METs) [71], one indoors (living room) and one outdoors (immediately outside the house) at each of four households per sub-village, operated from 18:00–07:00 (≈13 h), yielding 8 MET trap-nights per sub-village per sampling night; (ii) Backpack aspirator collections [72], conducted the following morning for 1 hour per house (4 hours of aspiration effort per site); and (iii) Barrier Screen Interception Traps (BS) [71], two screens per site operated from 18:00–07:00 (≈13 h), with collections conducted three times per night at standardized time windows: early evening (18:00–22:00), midnight (22:00–04:00), and pre-dawn (04:00–07:00), with approximately one hour of sampling within each window. A detailed description of sampling methods and sampling frequencies are found in the previous study [36].

### Morphological and molecular species identification

After collection, mosquitoes were morphologically identified at a complex/group or species level based on the protocols of Coetzee (2020) and Gillies and Coetzee (1987) [3,73] and stored individually in Eppendorf tubes filled with silica gel. Molecular species identification was performed using two approaches: species-specific PCR for a subset of specimens (n=2,642) morphologically identified as *An. gambiae* s.l. or *An. funestus* s.l. [39,40,74], while the larger subset of *Anopheles* samples (n=6,650) were preserved in 100% ethanol in 96-well plates and analysed using the ANOSPP amplicon sequencing panel [70]. For the ANOSPP dataset, DNA extraction was carried out using a minimally morphologically destructive protocol [75], and all samples went through the two-step ANOSPP PCR protocol and sequencing on Illumina MiSeq [70]. Species assignment was completed using the NNoVAE species assignment pipeline using *anospp_analysis v0.3.5* and reference datasets *nnv2* and *gcrefv1* [68,69].

### Post-NNoVAE species assignment of unresolved Funestus subgroup clades

Members of the *An. funestus* subgroup unresolved by the standard NNoVAE pipeline [69] were subjected to additional post hoc classification. These comprised Funestus_subgroup_cl1§ (*An. funestus* s.s., *An. funestus*-like, *An. vaneedeni*) and Funestus_subgroup_cl2§ (*An. parensis*, *An. longipalpis* C). PCA coordinate matrices were computed from 8-mer count tables concatenated across ANOSPP targets, using a minor-count frequency threshold of 0.02 with PCA implemented in scikit-learn v1.7.2 [76]. Reference specimens of known identity were used to identify species-informative axes in the PCA-reduced 8-mer space. Because leading PCs may capture broad genetic or population-structure variation rather than species-diagnostic separation, candidate PCs were selected using reference-only PC-wise one-way ANOVA with Benjamini–Hochberg correction. The selected PCs were then treated as the species-discriminating feature space. PERMANOVA tested overall multivariate separation among reference species and betadisper tested whether separation was attributable to unequal within-species dispersion (vegan::adonis2 and betadisper; 9,999 permutations) [77]. Final species assignment was performed in this selected PC space using a four-model bootstrap ensemble comprising k-nearest neighbours [78], radial-kernel support vector machine [79], random forest [80], and regularised quadratic discriminant analysis [81], representing complementary local, nonlinear, tree-based, and covariance-based classification rules. Models were trained across 2,000 balanced bootstrap resamples of the reference panel, and votes were aggregated across models and replicates to estimate support for each candidate species. Calls with ensemble support ≥0.80 were accepted as resolved, whereas lower-support calls were treated as unresolved and retained as *Anopheles funestus* s.l.

### *Plasmodium* species identification and phylogenetic tree generation

*Plasmodium* species identification is embedded into the ANOSPP protocol [70] through targeting two short mitochondrial amplicons across the whole genus (hereafter P1 and P2; ∼170–220 bp). Parasite presence was inferred from the recovery of *Plasmodium* reads, and primary species assignment for *Plasmodium*-positive samples was obtained using the ANOSPP species-assignment pipeline (*anospp_analysis v0.3.5*), which implements a local BLAST-based workflow against the *plasmv1* reference database [70]. To visualise sequence relationships and provide an additional layer of confirmation for these BLAST-based calls (BLAST+ 2.17.0), we constructed a local reference database of complete *Plasmodium* mitochondrial genomes by downloading sequences from NCBI via a custom Bash pipeline using E-utilities (*efetch,* EDirect 24.7). The combined FASTA file was organised by species and accession using a modified header format (>accession|species) and integrated into R (v4.5.2) workflows for phylogenetic analysis. Per-target alignments (sample amplicons plus matched reference segments) were generated with MAFFT v7.526 using the L-INS-i algorithm [82], and maximum-likelihood trees were inferred with IQ-TREE 3 v3.0.1 [83], employing ModelFinder for model selection and UFBoot (1,000 replicates) together with SH-aLRT (1,000 replicates) for branch support. Trees were rooted with *Haemoproteus columbae* as the outgroup and used qualitatively to verify that sample tips clustered with the reference sequences corresponding to their ANOSPP-assigned species (S2 Fig); *Plasmodium* species assignments reported in the main analyses are those produced by the ANOSPP pipeline.

### Statistical analyses

To evaluate the accuracy of morphological identification, ANOSPP assignments were treated as the reference standard. Concordance and misidentification rates were calculated for each morphologically defined taxon, with 95% binomial confidence intervals computed using the Wilson score method. Differences in misclassification frequency among taxa were assessed using a chi-squared test of independence; when expected cell counts were small, Fisher’s exact test was used (Monte Carlo simulation when required). Binomial logistic regression was used to estimate odds ratios (ORs) for misidentification relative to *An. coustani* s.l. as the reference category. Taxonomic diversity across collection sites was summarised using species richness and Shannon diversity (H′), calculated with the vegan package [84] in R (v4.2.2). Spatial patterns in species composition and diversity were visualised using ggplot2 [85], and maps were produced using Natural Earth shapefiles accessed via rnaturalearth [86].

District-level associations between primary-vector assemblages and malaria transmission intensity were evaluated using NMCP-defined transmission strata and the continuous NMCP composite score [64]. For each district, *An. arabiensis*, *An. funestus* s.s. and *An. gambiae* s.s. were coded as present or absent, with presence defined as at least one confirmed specimen of that species recorded in the district. These three binary variables were combined into an eight-level primary-vector assemblage variable representing all possible detection combinations: no primary vector detected, each single species, each two-species combination, and co-occurrence of all three species. Because this assemblage variable is categorical, its association with ordinal NMCP transmission stratum was tested using Fisher’s exact test on the observed contingency table, with p-values estimated by Monte Carlo simulation using 100,000 replicates. Primary-vector richness was then calculated as the number of primary-vector species detected per district, ranging from 0 to 3. Its association with the continuous NMCP composite score was tested using Spearman rank correlation, using the continuous score rather than the four ordinal strata to preserve variation along the transmission gradient. To assess whether this relationship was driven by broader differences in mosquito catch volume across districts, we also computed a partial Spearman correlation between primary-vector richness and the NMCP composite score while controlling for total *Anopheles* catch, defined as the total number of specimens assigned to all retained *Anopheles* taxa per district. Benjamini–Hochberg false-discovery-rate correction was applied across the three tests reported in this section: the Fisher’s exact assemblage test, the Spearman correlation between primary-vector richness and NMCP composite score, and the partial Spearman correlation controlling for total *Anopheles* catch.

## Results

### Mosquito morphological identification composition

Over a three-year surveillance period from December 2020 to December 2023, a total of 71,146 mosquitoes were collected across 25 sentinel districts. Morphological identification categorized these into two primary groups: culicines and anophelines (Fig 2). The majority of the collection, n = 61,126 (86.0%), were culicines. Within this group, species from the *Culex* genus predominated, accounting for n = 57,620 (80.99%) of the total collection. Other culicine genera identified included *Coquillettidia* at n = 1,942 (2.73%), *Mansonia* at n = 1,351 (1.90%), and *Aedes* at n = 213 (0.30%). The remaining n = 10,020 (14.0%) consisted of *Anopheles* species. Among these *Anopheles*, *An. gambiae* s.l. was the most prevalent, representing n = 4,168 (41.60%) of the *Anopheles* collection. Other notable *Anopheles* species included *An. pharoensis* at n = 2,005 (20.01%), *An. coustani* at n = 1,574 (15.71%), and *An. funestus* s.l. at n = 1,314 (13.11%). Additional *Anopheles* species identified in smaller numbers were *An. squamosus* (n = 905, 9.03%), *An. maculipalpis* (n = 18, 0.18%), *An. ziemanni* (n = 18, 0.18%), *An. rufipes* (n = 17, 0.17%), and *An. cinctus* (n = 1, 0.01%); this specimen was subsequently resolved by ANOSPP only to the *Christya* series due to limited reference sequences for this taxon (see ANOSPP-based Identification below).

**Fig 2.**
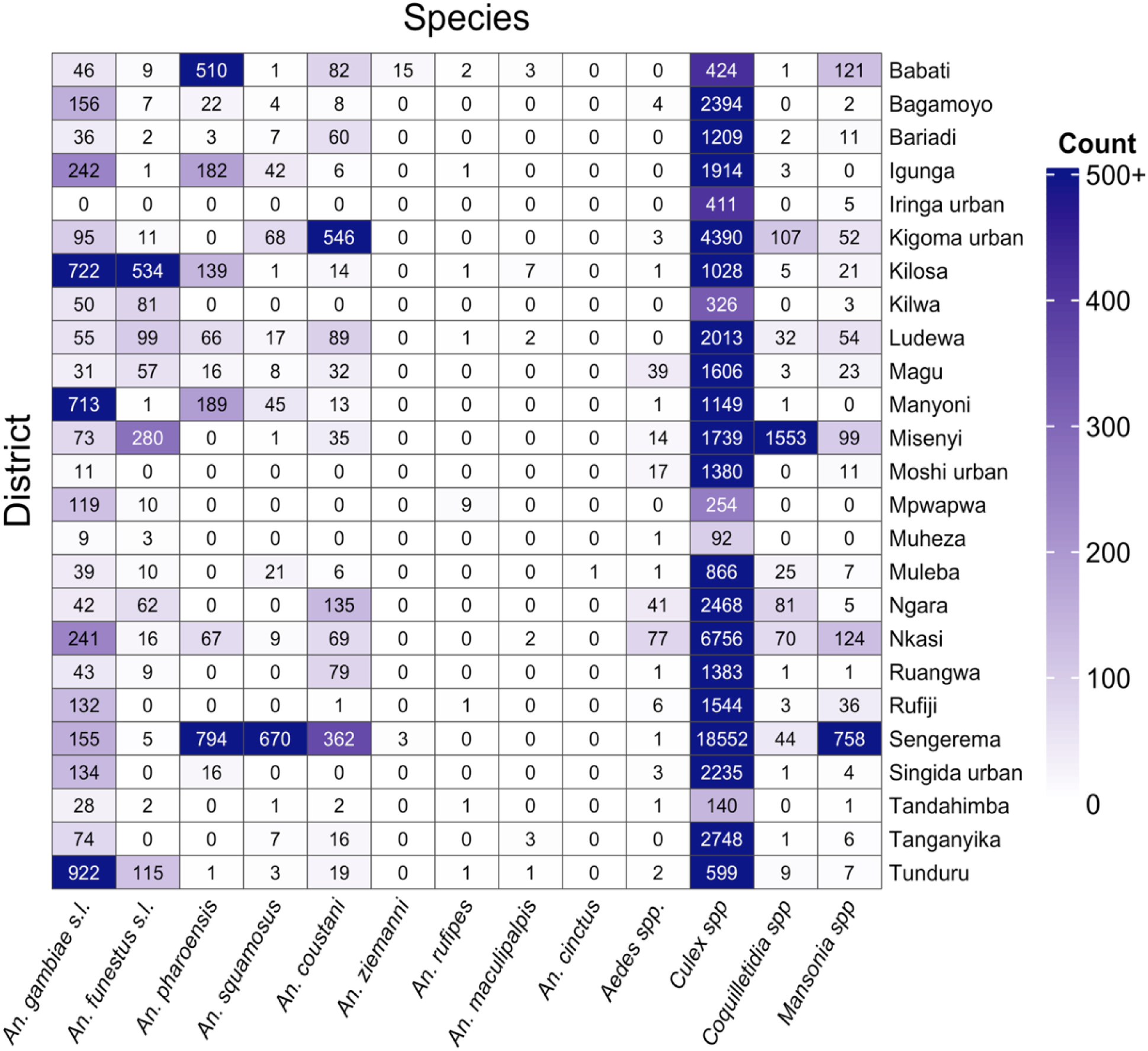
Distribution and total catches of mosquito taxa across districts as identified by morphological keys. The heatmap displays specimen counts by taxon (columns) and district (rows) collected during the study period. Cell colour intensity represents specimen count, with the gradient capped at 500 (counts above 500 shown at maximum saturation). Numeric values indicate the exact count per taxon–district combination.

### PCR-based identification

A total of 2,642 *Anopheles* mosquitoes, initially identified morphologically as members of the *An. gambiae* s.l. (2,096 individuals) or the *An. funestus* s.l. (546 individuals), were submitted for PCR amplification. Of these, 248 (9.39%) samples failed to amplify, yielding 2,394 (90.61%) successfully amplified and confirmed specimens. PCR-based identification revealed the following sibling species composition within the *An. gambiae* s.l. and *An. funestus* s.l.: *An. arabiensis* (n = 1,686; 70.42%), *An. gambiae* s.s. (n = 173; 7.23%), *An. quadriannulatus* (n = 62; 2.59%), *An. funestus* s.s. (n = 405; 16.92%), *An. leesoni* (n = 50; 2.09%), and *An. rivulorum* (n = 18; 0.75%). These specimens were not processed further using the ANOSPP panel, as PCR had already resolved them to species. Their PCR-based identifications were treated as final and pooled with ANOSPP-derived calls in subsequent analyses, except in the direct comparison of morphological and ANOSPP identifications, which was restricted to ANOSPP-typed specimens.

### ANOSPP-based identification

A total of 6,650 *Anopheles* mosquitoes were submitted for ANOSPP sequencing. Of these, 1,124 samples failed quality control due to insufficient amplicon recovery, likely attributable to poor sample quality, and 28 samples were further filtered out due to potential contamination. The remaining 5,498 samples were successfully identified at different levels of taxonomic resolution. The taxonomic resolution achieved across these samples varied. A small fraction (n = 10; 0.18%) were identified only at a coarse taxonomic level, either series or subgenus, indicating incomplete resolution: *Cellia* series (n = 3), *Christya* series (n = 1, comprising the single specimen morphologically identified as *An. cinctus*), and *Myzomyia* series (n = 6). An additional 999 samples (18.17%) were resolved at an intermediate level, typically complex or group level, also reflecting partial resolution: *An. coustani* s.l. (n = 669), *An. marshallii* s.l. (n = 41), *An. funestus* s.l. (n = 10), and *An. gambiae* s.l. (n = 279). Most samples were successfully resolved to the fine species level (n = 4,489; 81.65%), comprising 14 molecularly identified *Anopheles* taxa (Table 1). The resolved assemblage was dominated by two species with more than 1,000 individuals each: *An. pharoensis* (n = 1,573; 35.04%) and *An. arabiensis* (n = 1,361; 30.32%), which together accounted for 65.36% of fine-level assignments. Other relatively common taxa included *An. squamosus* (n = 671; 14.95%), *An. funestus* (n = 437; 9.73%), *An. rivulorum* (n = 170; 3.79%), and *An. ziemanni* (n = 121; 2.70%). The remaining taxa, including *An. gambiae* s.s., *An. quadriannulatus*, *An. pretoriensis*, *An. funestus*-like, *An. longipalpis* C, *An. parensis*, *An. rufipes*, and *An. maculipalpis*, each contributed less than 2% of fine-level assignments (Table 1).

**Table 1.** Species composition summary across *Anopheles* species complexes identified in this study.

| Series | Group/<br>Complex | Taxon<br>(molecular<br>assignment) | Vectorial status | PCR<br>(n) | ANOSPP<br>(n) | Total<br>(n) |
| --- | --- | --- | --- | --- | --- | --- |
| *Pyrethophorus* | *An. gambiae* s.l. | *An. gambiae* s.s. | Primary vector | 173 | 70 | 243 |
| *Pyrethophorus* | *An. gambiae* s.l. | *An. arabiensis* | Primary vector | 1,686 | 1,361 | 3,047 |
| *Pyrethophorus* | *An. gambiae* s.l. | *An. quadriannulatus* | Non-vector | 62 | 32 | 94 |
| *Pyrethophorus* | *An. gambiae* s.l. | Unresolved within<br>*An. gambiae* s.l. | Unresolved | - | 279 | 279 |
| *Myzomyia* | *An. funestus* s.l. | *An. funestus* s.s. | Primary vector | 405 | 437 | 842 |
| *Myzomyia* | *An. funestus* s.l. | *An. lesoni* | Secondary vector | 50 | 0 | 50 |
| *Myzomyia* | *An. funestus* s.l. | *An. rivulorum* | Secondary vector | 18 | 170 | 188 |
| *Myzomyia* | *An. funestus* s.l. | *An. longipalpis* C | Secondary vector | - | 13 | 13 |
| *Myzomyia* | *An. funestus* s.l. | *An. parensis* | Secondary vector | 0 | 6 | 6 |
| *Myzomyia* | *An. funestus* s.l. | *An. funestus*-like | Uncertain vector | - | 10 | 10 |
| *Myzomyia* | *An. funestus* s.l. | Unresolved within<br>*An. funestus* s.l. | Unresolved | - | 10 | 10 |
| *Myzomyia* | *An. marshallii* s.l. | Unresolved within<br>*An. marshallii* s.l. | Unresolved | - | 41 | 41 |
| *Myzomyia* | – | Unresolved within<br>*Myzomyia* series | Unresolved | - | 6 | 6 |
| *Cellia* | – | *An. pharoensis* | Secondary vector | - | 1,573 | 1,573 |
| *Cellia* | *An. squamosus* s.l. | *An. squamosus* | Secondary vector | - | 671 | 671 |
| *Cellia* | – | Unresolved within<br>*Cellia* series | Unresolved | - | 3 | 3 |
| *Christya* | – | Unresolved within<br>*Christya* series | Non-vector | - | 1 | 1 |
| *Neocellia* | – | *An. maculipalpis* | Non-vector | - | 3 | 3 |
| *Neocellia* | – | *An. pretoriensis* | Non-vector | - | 18 | 18 |
| *Neocellia* | – | *An. rufipes* | Secondary vector | - | 4 | 4 |
| *Myzorhynchus* | *An. coustani* s.l. | *An. ziemanni* | Secondary vector | - | 121 | 121 |
| *Myzorhynchus* | *An. coustani* s.l. | Unresolved within<br>*An. coustani* s.l. | Unresolved | - | 669 | 669 |
| **Grand total** |  |  |  | **2,394** | **5,498** | **7,892** |

**Note:** PCR and ANOSPP counts are based on different specimen subsets and should be interpreted within method-specific denominators. PCR counts represent successfully amplified specimens from the PCR-submitted subset (**n = 2,394/2,642**), which targeted mosquitoes morphologically assigned to *An. gambiae* s.l. and *An. funestus* s.l. ANOSPP counts represent retained assignments after sequencing quality control and contamination filtering (**n = 5,498/6,650**), including specimens resolved to species level (**n = 4,489**) and specimens assigned only to series, complex, or group level (**n = 1009**). Method-specific percentages are reported in the text. Dashes indicate taxa not targeted by the PCR workflow, rather than confirmed absence.

### Comparison of morphological and ANOSPP identification

Of the 5,498 specimens examined by ANOSPP, results indicated that 338 (6.2%) had been initially misclassified by morphological identification (95% CI: 5.5–6.8). Misidentification by morphology varied by taxon (χ² = 381.8, p = 0.001), with the lowest rate in *An. pharoensis* (0.6%) and the highest in *An. rufipes* (81.8%) and *An. maculipalpis* (77.8%) (Fig 3A). Notably, ANOSPP identified 41 specimens belonging to the *An. marshallii* s.l. and 18 specimens of *An. pretoriensis*, none of which had been distinguished as such in the original morphological sorting. Odds ratios showed the same pattern of relative misidentification risk (Fig 3B).

**Fig 3.**
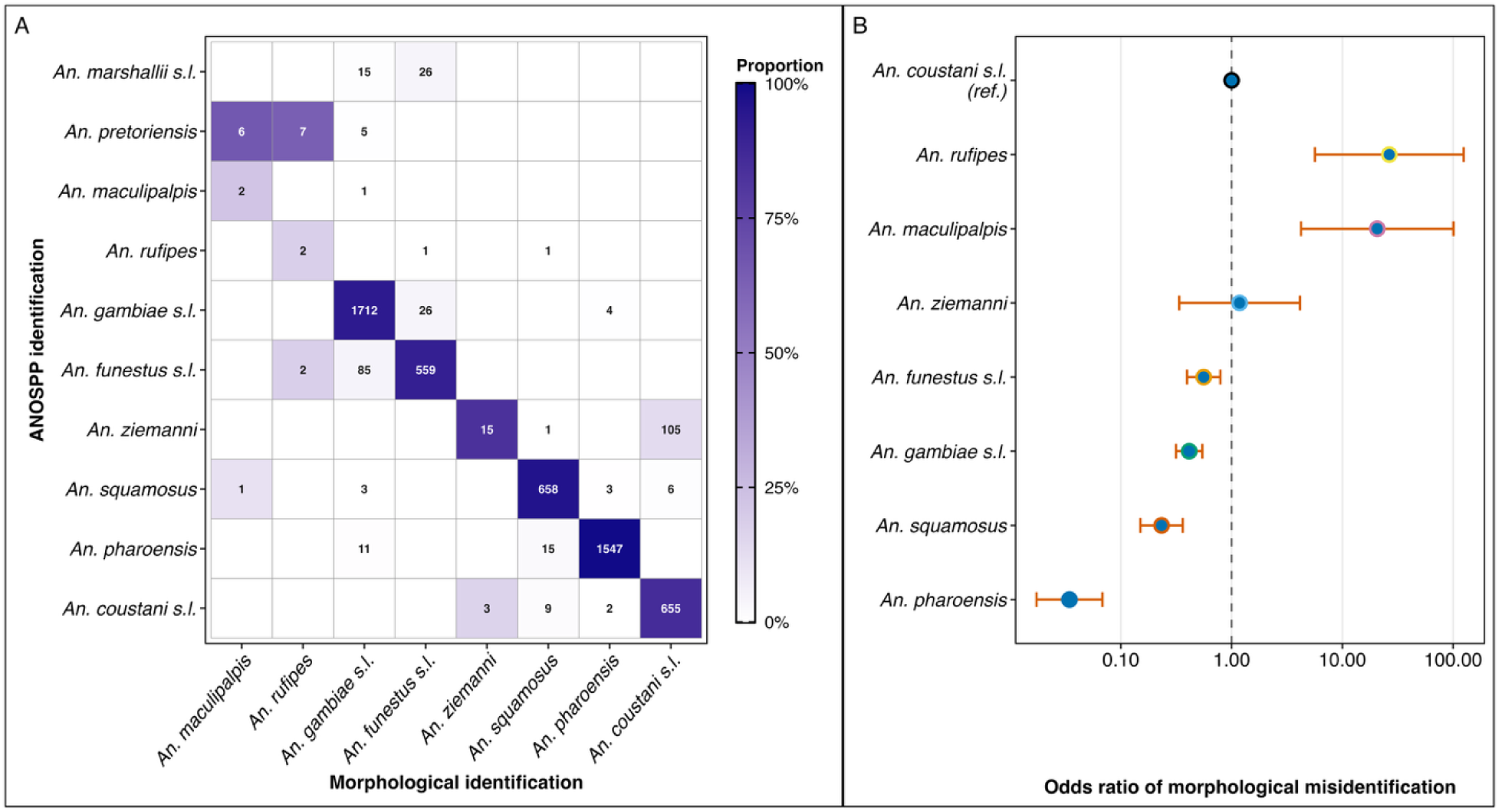
Concordance and misidentification patterns between morphological and ANOSPP-based species identification. **(A)** Heatmap of morphological versus ANOSPP identification. Each cell represents the proportion, shown by colour intensity, and the absolute number, shown as the numeric label, of specimens within each morphological group (columns) assigned to a given molecularly confirmed group by ANOSPP (rows). The proportion is calculated relative to the total number identified morphologically as each taxon. Diagonal cells indicate congruent identifications for all species except *An. marshallii* s.l. and *An. pretoriensis*, which were not detected morphologically; off-diagonal cells reveal misclassification patterns. **(B)** Forest plot showing odds ratios of morphological misidentification for each taxon relative to *An. coustani* s.l. as the reference group. Points denote odds ratio estimates, and horizontal bars show 95% confidence intervals. The vertical dashed line marks OR = 1, indicating no difference from the reference. Values above 1 indicate higher odds of misidentification than the reference, whereas values below 1 indicate lower odds. The x-axis is displayed on a logarithmic scale.

### *Anopheles* mosquito species spatial distribution and diversity

Figure 4A shows spatial distribution of the primary vector species across the country. *Anopheles arabiensis* was the most widely distributed primary vector, detected in 24 of 25 districts and contributing the largest proportional catches overall. In contrast, *An. gambiae* s.s. and *An. Funestus* s.s. showed more restricted distributions, occurring predominantly in the northwestern and southern regions of the country and being nearly absent from the relatively arid central corridor (S1 Fig). Of the secondary malaria vector species, *An. pharoensis* and *An. squamosus* were broadly distributed but showed relatively high catches in Sengerema, while members of the *An. Coustani* s.l. were similarly widespread with relatively high catches in Kigoma Urban (Fig 4B). The remaining species were generally more localized in distribution (Fig 4B). *Anopheles funestus*-like, the first confirmed record of this taxon in Tanzania was detected in Ludewa and Kilosa, while *An. longipalpis* Type C, also the first record of this cryptic type from Tanzania, was detected in two districts: Babati and Nkasi (Fig 4B). *Anopheles* species richness and diversity varied markedly between districts (Fig 4C, D), with three districts: Ludewa, Tunduru, and Kilosa consistently characterized by the highest richness and diversity, whereas Moshi Urban was characterized by the lowest richness and diversity. The district-level catches of each molecularly identified *Anopheles* species are provided in S1 Table.

**Fig 4.**
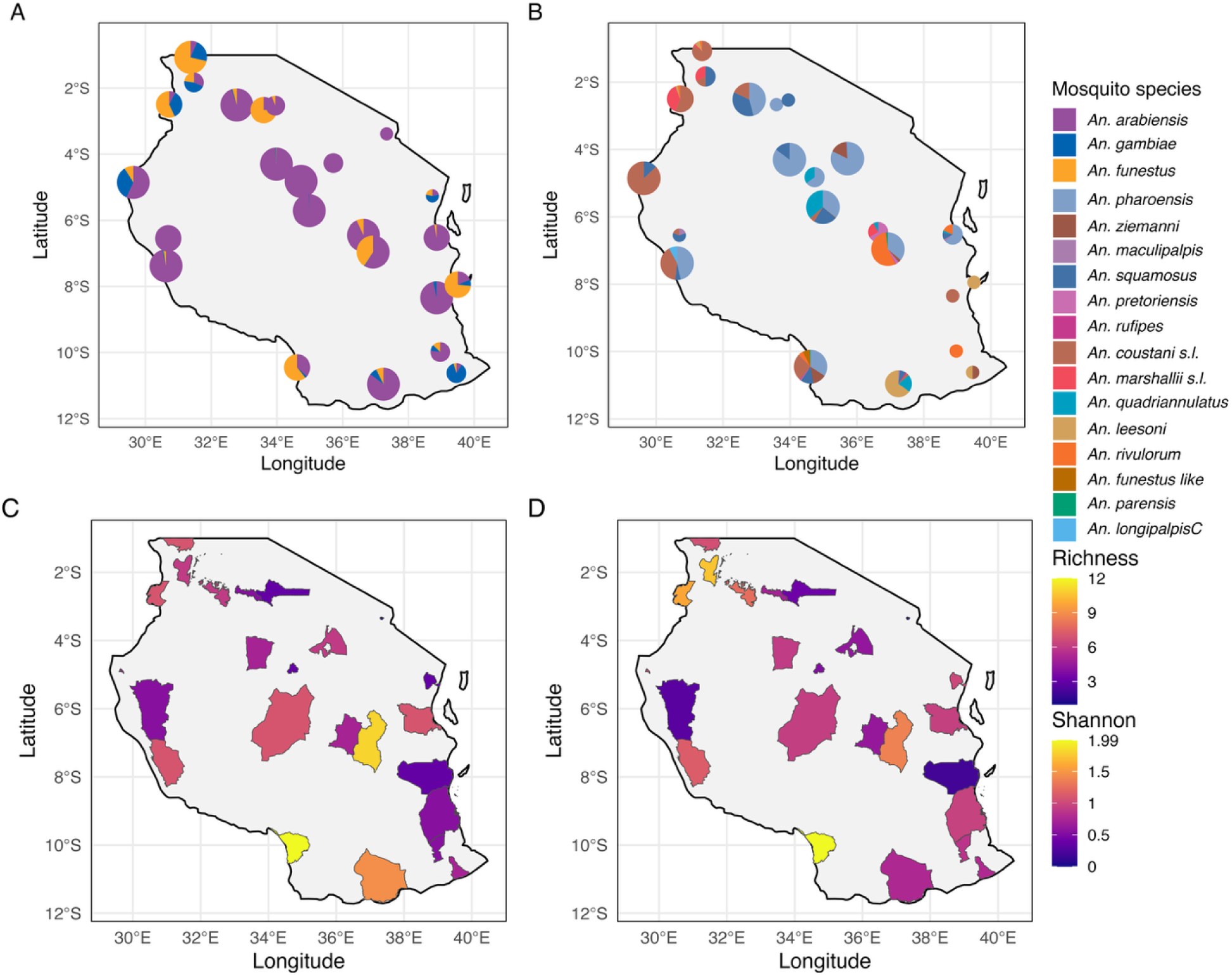
Geographic Patterns of *Anopheles* Species Composition, Richness, and Diversity. Panel **A** displays the composition of primary malaria vectors (*An. arabiensis*, *An. gambiae* s.s., and *An. funestus* s.s.), while panel **B** shows secondary vectors and other *Anopheles* species. Pie charts show the proportional composition of mosquito species at each site. Pie size is scaled to the total number of mosquitoes sampled per district, with larger pies indicating higher catches (categories: <10, <50, <100, ≥100). Panel **C** illustrates species richness (total number of distinct *Anopheles* species), while Panel **D** shows the Shannon diversity index (H’), which accounts for both species richness and the relative evenness of species catches across districts. Higher values in both metrics indicate greater ecological complexity of mosquito populations across regions.

### The distribution of primary vectors aligns with national malaria transmission strata

District-level combinations of *An. arabiensis*, *An. funestus* s.s., and *An. gambiae* s.s. differed significantly across NMCP transmission strata (Fisher’s exact test, BH-adjusted *p* = 0.006). Very-low and low-transmission strata were mainly characterised by *An. arabiensis* alone or by the absence of any detected primary vector species, whereas moderate- and high-transmission strata increasingly exhibited multiple primary vectors, with three-species combinations (*An. arabiensis* + *An. funestus* s.s. + *An. gambiae* s.s.) dominating most high-transmission strata. The number of primary-vector species detected per district was strongly correlated with the NMCP composite transmission score (Spearman ρ = 0.722, BH-adjusted *p* < 0.001), and this association remained robust after partialling out total *Anopheles* catch (partial ρ = 0.765, BH-adjusted *p* < 0.001), indicating that the richness signal was independent of overall catch volume.

### *Plasmodium* species detected in *Anopheles* mosquitoes and their spatial distribution

Five *Plasmodium* species were detected across five *Anopheles* taxa (Table 2). *Anopheles funestus* s.s. carried the broadest parasite diversity (*P. falciparum*, *P. malariae*, *P. ovale*, and *P. vivax*) and the largest absolute number of positive specimens (20/437; 4.58%). It was also the only vector species in which mixed-species *Plasmodium* detections occurred in individual mosquitoes all from Ngara District. *Anopheles gambiae* s.s. was positive only for *P. falciparum* but showed the highest point estimate of positivity (4/70; 5.71%), while *An. arabiensis* had detections of *P. falciparum* (5/1361; 0.37%) and *P. caprae* (6/1361; 0.44%). The detection of *P. caprae*, a goat-associated ungulate malaria parasite, in Kilosa district, represents both the first record of *P. caprae* in Tanzania and the first detection of this parasite in *An. arabiensis* globally. Single *P. falciparum* detections were also recorded in *An. pharoensis* (1/1,573; 0.06%) and *An. rivulorum* (1/170; 0.59%). Across parasites, *P. falciparum* was the most frequently detected and most widely distributed, whereas the other *Plasmodium* species showed more restricted distributions (Fig 5). Phylogenetic placement of P1 and P2 amplicons was concordant with ANOSPP-assigned *Plasmodium* species for all reported positive specimens (S2 Fig).

**Fig 5.**
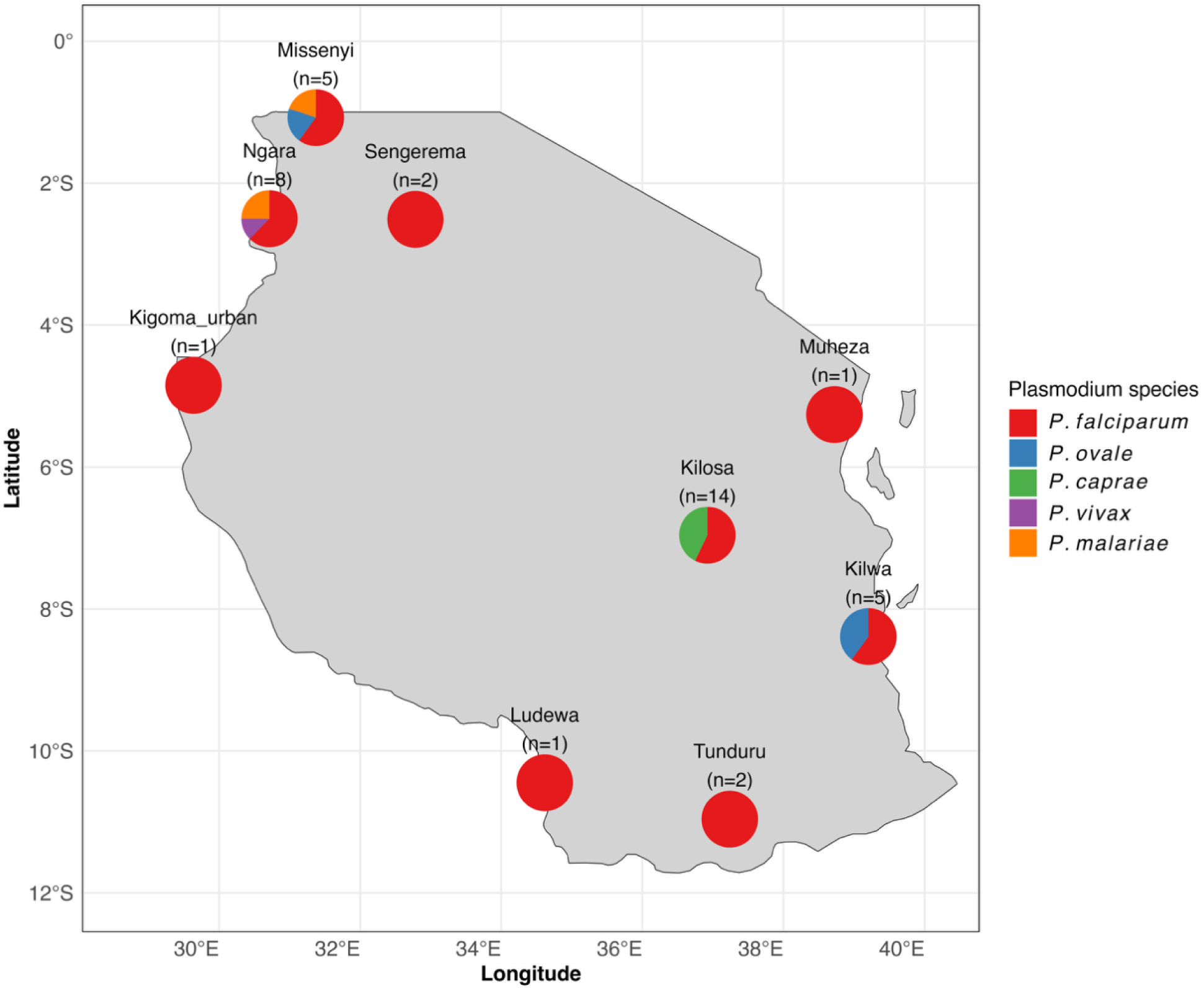
Spatial distribution of *Plasmodium* detected in mosquito vectors across surveyed districts. Each district pie chart shows the proportional composition of the five *Plasmodium* species identified. District counts (*n*) reflect parasite detections; two mixed-species infections in *An. funestus* s.s. (Ngara) are counted once per species.

**Table 2.** Detection Status of Individual *Anopheles* Mosquitoes Harbouring *Plasmodium* Parasites.

| **Mosquitoes Species** | ***Plasmodium* Status** | ***Plasmodium* Species** | **Count** | **Prevalence (Numbers)** | **Prevalence (Percent)** |
| --- | --- | --- | --- | --- | --- |
| *An. arabiensis* | Single species | *P. falciparum* | 5 | 5/1361 | 0.37 |
| *An. arabiensis* | Single species | *P. caprae* | 6 | 6/1361 | 0.44 |
| *An. funestus* | Two species | *P. vivax*; *P. malariae* | 1 | 1/437 | 0.23 |
| *An. funestus* | Two species | *P. falciparum*; *P. malariae* | 1 | 1/437 | 0.23 |
| *An. funestus* | Single species | *P. falciparum* | 14 | 14/437 | 3.20 |
| *An. funestus* | Single species | *P. malariae* | 1 | 1/437 | 0.23 |
| *An. funestus* | Single species | *P. ovale* | 3 | 3/437 | 0.69 |
| *An. gambiae* | Single species | *P. falciparum* | 4 | 4/70 | 5.71 |
| *An. pharoensis* | Single species | *P. falciparum* | 1 | 1/ 1573 | 0.06 |
| *An. rivulorum* | Single species | *P. falciparum* | 1 | 1/170 | 0.59 |

## Discussion

Accurate identification of malaria vectors remains a critical bottleneck for epidemiological inference and for designing species-appropriate control interventions. In this nationwide survey, we show that standard morphological identification, although widely used in Tanzania, still misclassifies a non-trivial fraction of specimens (6.2%) when benchmarked against ANOSPP-based assignments, with substantially higher misclassification for rare taxa. Although this overall misclassification rate is lower than those reported in some neighbouring settings, for instance 15% in Zambia [87], and 10.8% [88] and 16% [89] in Kenya, both benchmarked against ITS2 and COI sequencing, the aggregate error conceals important taxonomic skew. Errors are not evenly distributed: infrequent or morphologically challenging species are disproportionately affected [11,43,44,90]. In our data, *An. rufipes* and *An. maculipalpis* were frequently misidentified, while *An. pretoriensis* and members of *An. marshallii* s.l. were never recognised morphologically and would have been completely overlooked without ANOSPP. These misidentifications matter because species within *An. marshallii* s.l. have been found carrying sporozoites [17,19,91,92], while *An. rufipes*, which has historically been ignored, is increasingly recognised as an important vector [87,93,94]. Although these taxa were found in low numbers in our survey and none tested positive for *Plasmodium*, their misclassification would still remove potentially relevant secondary vectors from the surveillance picture. Thus, even a well-designed nationwide survey relying on morphology alone would have produced an incomplete picture of anopheline diversity and of the taxa potentially relevant to local malaria transmission.

Beyond the disproportionate misidentification of rare or non-primary taxa, morphological keys cannot reliably resolve species within the *An. gambiae* complex and the *An. funestus* group, where morphological characteristics are often identical [20]. In practice, this limitation propagates into downstream molecular workflows: many widely used ITS2-based PCR assays require correct group-level identification, target only a subset of species, and are vulnerable to primer-binding polymorphisms [11], so early mislabelling or sequence variation in primer-binding sites can result in non-amplification or spurious assignments [20,41]. Because morphologically similar species can differ in behaviour, ecology, and vectorial capacity, such errors can distort inference on which vector groups are present, where they occur, and how interventions should be evaluated [28,42]. In our survey, for instance, some ANOSPP-assigned non-primary taxa were recorded morphologically as major vector groups: members of *An. marshallii* s.l. were labelled as *An. Gambiae* s.l. or *An. funestus* s.l., while *An. pretoriensis* and *An. pharoensis* were recorded as *An. gambiae* s.l. Conversely, some ANOSPP-assigned *An. funestus* s.l. specimens were sometimes morphologically labelled as *An. gambiae* s.l., and vice versa. Such errors would not only affect taxonomic lists; they could distort estimates of the relative distribution and abundance of the vector groups most often used to guide intervention planning and evaluation. By contrast, ANOSPP leverages 62 loci distributed across the mosquito genome and uses k-mer-based assignment against curated reference sets, making species calls more robust to local variation at any single locus and reducing dependence on detailed morphological pre-sorting [68–70]. In our study, this translated into higher overall taxonomic resolution and systematic recovery of several routinely neglected taxa, including new country records and the first country-level mapping of multiple secondary vectors using a single, standardised assay.

This expanded molecular resolution enhanced our view of anopheline diversity in Tanzania, revealing 15 *Anopheles* species and providing a contemporary geo-referenced account of their distribution. Notably, we detected *An. funestus-like*, principally from Manda Ward, Ludewa District, representing, to our knowledge, the first confirmed Tanzanian record. This extends its documented range from northern Malawi into southern Tanzania and is biogeographically plausible given Manda’s proximity to Karonga Region, where this taxon has been recorded [95,96]. Most specimens were collected outdoors near livestock shelters, consistent with the predominantly zoophilic behaviour reported elsewhere [96]. Its additional detection in Kilosa District suggests that *An. funestus-like* may have a broader distribution within Tanzania. We also refined records of *An. longipalpis* by identifying *An. longipalpis* Type C in Nkasi and Babati districts. Although *An. longipalpis* has previously been reported from several Tanzanian localities [38], these records were largely morphology-based and not resolved into the provisionally designated Type A and Type C taxa recognised from molecular studies [97,98]. This distinction is important because forms historically assigned to *An. longipalpis* represent phylogenetically distinct lineages, with Type A associated with the *An. minimus* subgroup and Type C with the *An. funestus* subgroup [69,99,100]. They also differ in vector implication: Type C has been found resting indoors [98,101], and recent surveys in Kenya reported *P. falciparum* sporozoite infection [102,103], including a high outdoor sporozoite rate [103]. These findings implicate Type C as a potential contributor to residual malaria transmission in some settings [102,103]. *Anopheles longipalpis* Type A, by contrast, has not been implicated in *Plasmodium* transmission, and its ecology remains insufficiently characterised [98,104,105]. Detection of Type C therefore provides contemporary molecular evidence of its occurrence in Tanzania and warrants further studies to clarify its local transmission role. Beyond these headline findings, ANOSPP also confirmed rarely documented taxa, including *An. pretoriensis* [106,107] and *An. marshallii* s.l. [12,17,108].

For several species identified in this survey, documented distributions remain sparse, with many records being historical and morphology-based [38]. This gap is epidemiologically important because historically neglected taxa are increasingly implicated in local malaria transmission across African settings [43–45,93,94]. In Tanzania, the clearest contemporary example is *An. parensis*, recently confirmed as a secondary vector through a molecularly informed nationwide survey [11]. For other non-primary vectors, evidence remains more fragmentary, drawn largely from older records concentrated in present-day Tanga and Kilimanjaro regions, with limited systematic follow-up elsewhere [5,8,12,15–19,109]. Against this backdrop, our study strengthens the contemporary georeferenced evidence base for *Anopheles* in Tanzania, with 15 resolved species. Several additional taxa remain unresolved at subgroup, complex, or group level and could be fully resolved once additional reference sequences become available [69]. Together, these findings update the known distribution of *Anopheles* taxa in Tanzania by adding new country records, refining the species identity of historically morphology-based records, and providing contemporary molecular confirmation for several rarely documented taxa.

Across the *Anopheles* species, we detected five *Plasmodium* species, namely *P. falciparum*, *P. malariae*, *P. ovale*, *P. vivax* and *P. caprae*, the latter detected only in *An. arabiensis*. This represents, to our knowledge, the first record of *P. caprae* in Tanzania and its first detection in *An. arabiensis*. *Plasmodium caprae* is a goat malaria parasite previously reported from parts of Asia and Africa [110–112], including detections in *An. subpictus* and *An. aconitus* [111], but, to our knowledge, has not been reported in humans. Its detection in *An. arabiensis* may indicate either that this vector can carry a broader range of *Plasmodium* parasites than previously recognised, or that the finding reflects a recent goat blood meal due to its opportunistic feeding on domestic goats [21,36]. Detection of a goat malaria parasite in a recognised human malaria vector illustrates how surveillance frameworks focused exclusively on *P. falciparum* risk missing broader *Plasmodium* diversity at the human-animal interface [61].

Overall, *An. funestus* s.s. showed the broadest parasite diversity and accounted for the largest number of parasite-positive specimens, whereas *An. gambiae* s.s. had the highest point estimate of parasite positivity. Although detections in secondary vector taxa were sparse, their occurrence in *An. rivulorum* and *An. pharoensis* indicates that parasites were not confined to the major vector species [5,12,113]. Beyond *P. falciparum*, which is routinely targeted and therefore commonly detected in entomological surveillance across Tanzania [1,10,11], field detection of non-*falciparum* parasites in local vectors has rarely been reported. *Plasmodium ovale* has previously been demonstrated in *Anopheles* in Tanzania only under experimental skin-feeding conditions [32], and to our knowledge there are no prior records of *P. malariae* in wild-caught Tanzanian *Anopheles*, despite both species being well documented in human infections nationally [2,33,114]. Field evidence of *P. vivax* in African vectors remains rare [115,116], and to our knowledge this is the first molecularly confirmed detection in a wild-caught Tanzanian *Anopheles*, consistent with growing recognition of *P. vivax* transmission across sub-Saharan Africa [116]. Because ANOSPP detects *Plasmodium* DNA rather than stage-specific sporozoite infection, these findings provide evidence of local parasite circulation and, for human malaria parasites, are consistent with infections in the local human population, but they do not by themselves demonstrate established vector infection or onward transmissibility. Nonetheless, detecting multiple *Plasmodium* species across both primary and secondary vectors in a single assay is rarely achieved in routine surveillance, which predominantly focuses on *P. falciparum* [1]. Together with improved mosquito species assignment, this parasite-resolution capacity underscores the value of ANOSPP as an integrated tool for vector and parasite surveillance.

*Plasmodium falciparum* remained by far the dominant species detected. Several high-transmission districts, particularly Ngara, Missenyi and Kilwa [64], stood out as foci of broader *Plasmodium* species diversity. In these locations, *P. falciparum* co-occurred with *P. malariae*, *P. ovale* and/or *P. vivax*, suggesting that areas with higher malaria transmission may also sustain a wider range of human *Plasmodium* species. Such co-circulation can complicate case management, especially where routine diagnosis relies heavily on malaria rapid diagnostic tests (mRDTs), because mixed or non-*falciparum* infections may be under-resolved and different species have distinct clinical manifestations and relapse profiles [54,55,59,60,117,118]. These findings argue that even in settings where *P. falciparum* clearly dominates, non-*falciparum* species can add an additional layer of complexity that is easily missed by *falciparum*-focused surveillance and control strategies.

Notably, the composition of primary malaria vectors varied across national malaria transmission strata. Higher-transmission settings harboured multiple primary vectors, particularly *An. arabiensis*, *An. funestus* s.s., and *An. gambiae* s.s., whereas very-low and low-transmission settings were often characterised by the presence of *An. arabiensis* alone or by the absence of any detected primary vector. This suggests that, under contemporary Tanzanian ecological and intervention contexts, the realised contribution of *An. arabiensis* to malaria transmission may be more context-dependent than implied by its traditional designation as a primary vector, although direct measurement of species-specific transmission (e.g. entomological inoculation rate (EIR) across strata) would be needed to confirm this interpretation. Recent studies from Tanzania [36] and Kenya [89] indicate that *An. arabiensis* is predominantly bovine-fed and exhibits a very low human blood index. This could explain why previous entomological inoculation rate estimates have shown disproportionate transmission by *An. funestus* s.s. where it co-occurs with *An. arabiensis* [5,119,120], consistent with the Ross–Macdonald model of transmission [121,122], which identifies vector survival and human-feeding preference as the most sensitive determinants of transmission intensity, both of which favour *An. funestus* s.s., which feeds preferentially on humans [24] and survives longer than *An. arabiensis* [67].

This study was not without limitations. First, although the study covered 25 districts spanning all NMCP transmission strata, surveillance was restricted to NMCP sentinel sites and may not fully represent the national distribution of vectors and parasites. Second, some *Anopheles* specimens remained unresolved at species level because appropriate reference sequences were unavailable or amplicon recovery was insufficient. Continued expansion and curation of reference datasets will therefore be essential to improve the resolution power of ANOSPP. Third, the wider application of ANOSPP remains restricted because sequencing is currently conducted at the Wellcome Sanger Institute [68,69], limiting the timely processing of samples. Broader application of ANOSPP will depend on strengthening in-country sequencing and bioinformatics capacity, alongside targeted capacity strengthening and skills transfer initiatives.

Despite these limitations, this study provides a unique and comprehensive view of contemporary *Anopheles–Plasmodium* diversity and distribution in Tanzania. It shows that Tanzania’s malaria-vector landscape is more complex than conventional surveillance approaches reveal. Combining ANOSPP with conventional entomological methods substantially improves the resolution and reliability of malaria vector and parasite surveillance. As reference databases expand and sequencing capacity becomes more accessible within Tanzania and comparable settings, targeted panels such as ANOSPP could strengthen NMCP monitoring and help ensure that malaria control strategies remain aligned with the actual and evolving diversity of vectors and parasites sustaining transmission.

## Acknowledgments

We extend our sincere gratitude to the local leaders and residents across the twenty-five study districts for their collaboration and support throughout the research. We are especially grateful to all volunteers who generously participated in this study. We also thank Mr. Eldad Govella, our driver, for his unwavering dedication and logistical support that greatly facilitated field activities. We further acknowledge Mr. Dickson Msaky for his sustained technical support with electronic data capture, including iterative reformatting and maintenance of the then MosquitoDB system, which was instrumental in ensuring smooth field data collection.

## Supporting information

**S1 Fig. Climatic context and malaria transmission strata of study sites in mainland Tanzania.** (A) Sentinel districts coloured by National Malaria Control Programme malaria transmission strata (Very low, Low, Moderate, High), with sampling sites shown as white points; sentinel districts are labelled by black numbers. (B) Sampling sites (white points) overlaid on the Köppen–Geiger climate classification map of Tanzania. Climate classes shown are: Af (tropical rainforest), Am (tropical monsoon), Aw (tropical savanna), BWh (hot desert), BSh (hot semi-arid steppe), Cfa (humid subtropical), Cwb (temperate oceanic/subtropical highland with dry winters), Cwc (subtropical highland with dry winters), ET (tundra; near the summit of Mount Kilimanjaro), and EF (ice cap; summit of Mount Kilimanjaro). This figure is adapted from Fig 1 of Baravuga et al. (2026), *Malaria Journal* 25:207 [36], published under a Creative Commons Attribution 4.0 (CC BY 4.0) licence.

**S2 Fig. Maximum-likelihood phylogenetic trees for *Plasmodium* species identification.** Phylogenetic trees inferred from ANOSPP mitochondrial amplicons P1 (A) and P2 (B). Trees were reconstructed in IQ-TREE 3 under the best-fit nucleotide substitution models selected by ModelFinder. Node support values represent ultrafast bootstrap (UFBoot) and SH-aLRT estimates. Each tip corresponds to either a reference mitochondrial genome (circles) or a study-derived haplotype (triangles). Major clades are colour-coded by species according to the key on the right. Trees are rooted with *Haemoproteus columbae* as the outgroup, and branch lengths indicate substitutions per site.

**S1 Table. District-level malaria transmission strata and *Anopheles* species captured.** Each column shows the number of mosquitoes identified to a given *Anopheles* species in each district, stratified by national malaria transmission category (Very low, Low, Moderate, High). Members of the *An. coustani* and *An. marshallii* complexes were reported at group level (“*An. coustani* s.l.”, “*An. marshallii* s.l.”) where sequencing supported membership of the complex/series but did not allow finer resolution.

